# STUN: forward-time Simulation on TUnable fitNess landscapes in recombining populations

**DOI:** 10.1101/2023.02.15.528681

**Authors:** André Amado, Juan Li, Claudia Bank

## Abstract

Understanding the population genetics of complex polygenic traits during adaptation is challenging. Here, we implement a forward-in-time population-genetic simulator (STUN) based on Wright-Fisher dynamics. STUN is a flexible and user-friendly software package for simulating the polygenic adaptation of recombining haploid populations using either new mutations or standing genetic variation. STUN assumes that populations adapt to sudden environmental changes by undergoing selection on a new fitness landscape. With pre-implemented fitness landscape models like Rough Mount Fuji, NK, Block, additive, and House-of-Cards, users can explore the effect of different levels of epistasis (ruggedness of the fitness landscape). Custom fitness landscapes and recombination maps can also be defined. STUN empowers both experimentalists and advanced programmers to study the evolution of complex polygenic traits and to dissect the adaptation process.

**Availability and implementation:** STUN is implemented in Rust. Its source code is available at https://github.com/banklab/STUN, which also includes a link to the software’s manual and binary files for Linux, macOS and Windows. Upon publication, the code will also be archived on Zenodo and a doi will be provided here.

## 1 Introduction

Understanding the genetic basis and dynamics of complex traits poses a significant challenge in population and quantitative genetics (Walsh and Lynch 2018). When a population encounters a sudden environmental change, adaptation often occurs through changes in polygenic traits. Both empirical and theoretical studies have been conducted to investigate the genetic basis of such traits. However, identifying the specific genes responsible and understanding their adaptive consequences is challenging due to the vastness of the underlying genotype space. The unpredictable nature of evolution within this space makes it difficult to disentangle and reconstruct the exact adaptive path. In this context, computational approaches play a crucial role in exploring potential scenarios of adaptation. They provide a means to simulate and study the dynamics of complex genetic systems under new environmental conditions. However, there is a scarcity of simulators that efficiently and flexibly incorporate recombining populations adapting on complex fitness landscapes.

Existing simulators, such as simuPOP (Peng and Kimmel 2005), SLiM Haller and Messer 2019 and QuantiNemo (Neuenschwander et al. 2019), primarily focus on direct, non-epistatic selection and dominance at a single locus, without straightforward options for consideration of the underlying genetic architecture arising from epistatic interactions. Although QuantiNemo and SLiM provide the option to assign an arbitrary fitness value to a genotype and thus, in principle, allow for an association between genotypes and fitness, they do not provide a large-scale (or probabilistic) means to define or characterize the ruggedness of the fitness landscape in terms of the relative contributions of non-additive and additive genetic effects. Furthermore, when we consider non-additive effects (epistasis and dominance) on a fitness landscape, their interplay requires complicated assumptions and entails a large computational burden. Although epistasis is expected to be ubiquitous and potentially complex in nature (Bank 2022), we are not aware of any existing software package to comprehensively perform evolutionary simulations on fitness landscapes with varying levels of epistasis (tunable ruggedness). To bridge this gap, we present STUN, an easy-to-use and open-source simulator that enables customization of fitness landscapes, recombination, new mutations and standing genetic variation in the model and that captures the resulting complex adaptive process. Importantly, STUN goes beyond solely recording the population genotype composition by offering a range of statistics on population features. To support users in applying STUN effectively, STUN is accompanied by a comprehensive manual, which includes detailed explanations of fitness landscape models and hands-on tutorials to guide users through the practical use of the software package.

## 2 Results

### 2.1 Overview of STUN

STUN takes advantage of a diverse range of fitness landscape models, as described in Fragata et al. 2019, to simulate the adaptation of haploid populations with recombination on arbitrary fitness landscapes (Figure 1). The genetic variants in the simulated populations are derived either through mutations or standing genetic variation initialized based on the neutral site frequency spectrum. STUN follows a Wright-Fisher model, assuming non-overlapping and discrete generations. In each generation, individuals in the population are randomly paired, and recombination is performed to generate a pool of gametes. Subsequently, haploid offspring are sampled from this gamete pool with probabilities proportional to their fitness values. These offspring then form the new generation. This process continues until either all loci in the population become monomorphic or a custom-defined stopping criterion is met. By following this model, STUN allows for the simulation of adaptive processes, incorporating recombination and the complex selection effects of fitness landscapes. This enables researchers to study population-genetic dynamics in the context of complex genetic architectures and rugged fitness landscapes.

**Figure 1.**
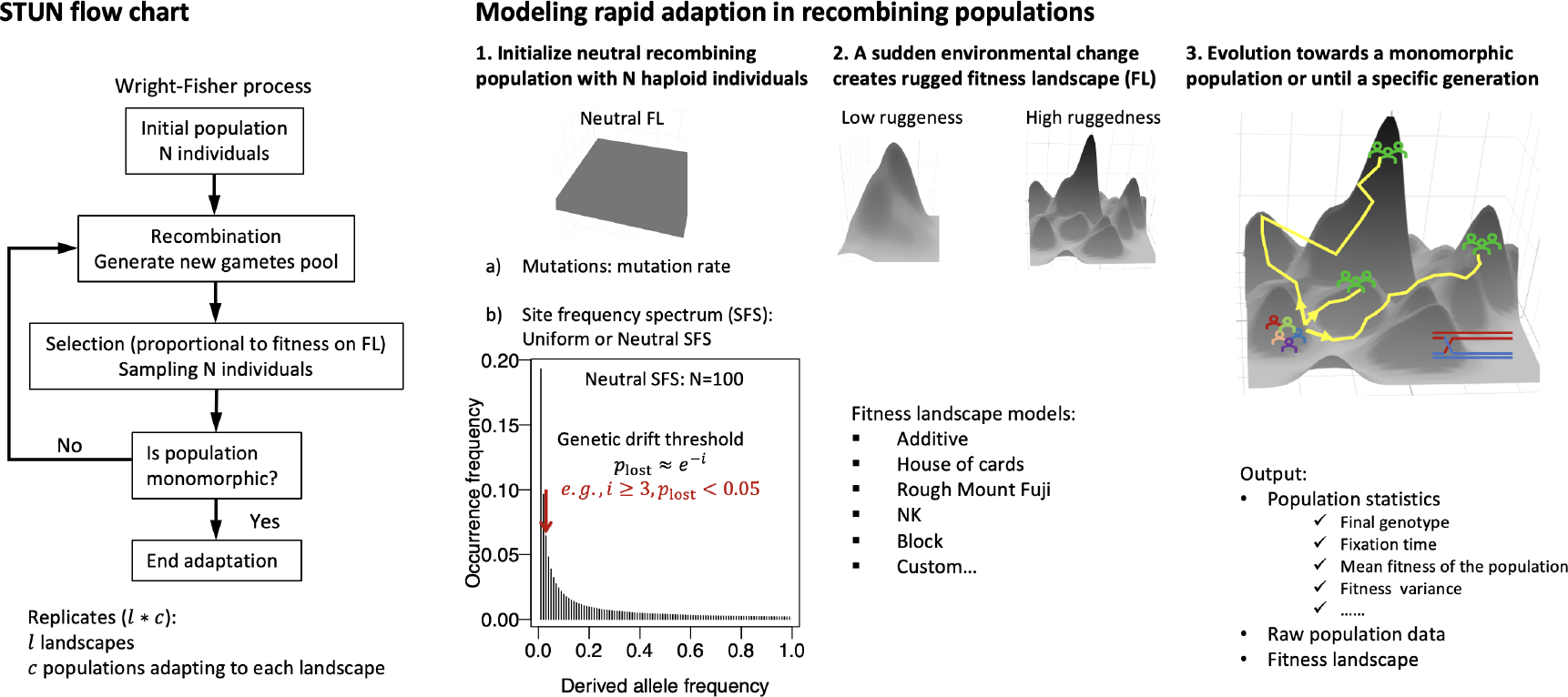
Overview of STUN. STUN is a population-genetic simulation package designed for studying complex trait adaptation. It employs Wright-Fisher dynamics and incorporates rugged fitness landscapes. STUN also offers a range of statistics on population features.

### 2.2 Features

#### Pre-defined and custom fitness landscapes

STUN includes a fitness landscape generator that offers out-of-the-box implementations of five commonly used fitness landscape models: the Rough Mount Fuji (Aita et al. 2000), NK (Kauffman and Levin 1987), Block (Perelson and Macken 1995), additive, and House-of-Cards models (reviewed in Fragata et al. 2019). The first three models allow for the generation of tunably rugged fitness landscapes. The ruggedness of a fitness landscape refers to the relative contribution of epistasis (genetic interactions between loci for fitness) compared to additive single-locus selection. The additive model assumes no epistasis, resulting in a ruggedness of 0 of the fitness landscape. This model serves as the no-epistasis limiting case for the Rough Mount Fuji model (by setting the epistatic parameter to zero), NK model (by setting the number of interacting loci *K* to zero), and Block model (by setting the number of blocks equal to the number of loci). On the other hand, the House-of-Cards model assigns statistically independent fitness values to each genotype, representing the maximum ruggedness limit of the Rough Mount Fuji model (by setting the additive contribution to zero), NK model (by setting the number of interacting loci *K* to *N −* 1), or Block model (by setting the number of blocks to 1). In addition to these pre-defined models, STUN also supports the specification of user-defined fitness landscapes, hereby, e.g., allowing for the simulation of adaptation on experimentally inferred fitness landscapes.

#### De novo mutations and standing genetic variations

STUN offers two sources of genetic variants: initial standing genetic variation and new mutations. In the case of standing genetic variation, a population with specified initial allele frequencies is generated initially. The allele frequencies at each locus can be equal or drawn from a folded neutral site frequency spectrum (SFS, Hudson 2015). To avoid the generation of low-frequency alleles that are not subject to significant selection during the early generations, a genetic drift threshold can be set for the neutral SFS (for details, see Li et al. 2023). This threshold considers the probability of an allele being lost in the first generation after initializing, which is calculated as *e*^*−i*^, where *i* represents the copy of alleles in the initial population. To ensure a low probability of initial random loss of an allele (less than 0.05), it is recommended to have at least three copies of each allele in a population of relatively large size (greater than 100). In addition, STUN allows for new mutations by assigning a mutation rate. With a low mutation rate, the classical assumption of strong-selection-weak-mutation is met (SSWM, Gillespie 1984), whereas a high mutation rates result in multiple mutations segregating simultaneously, providing the opportunity to capture interference effects that violate the assumption of SSWM and potentially alter the outcome of adaptation.

#### Recombination

The balance between selection and recombination has been of great interest when studying quantitative traits, even in the absence of epistasis, as observed in phenomena such as Hill-Robertson interference (Hill and Robertson 1966). It has been recognized that recombination can confer an advantage during the early stages of adaptation because epistasis relies on the linkage of mutations, thereby suggesting a temporary advantage of sexual reproduction (Nowak et al. 2014). This highlights the significance of individual-based simulations, such as those implemented in STUN, in providing valuable insights and predictions for complex traits that cannot be adequately addressed through analytical approaches alone.

Unlike the approach described in Nowak et al. 2014, which replaces a fraction of individuals with recombinants of two randomly chosen parents, STUN offers two realistic options for defining recombination. The first option is a uniform recombination map, where the recombination probability is assumed to be constant between neighboring loci. The second option allows for the specification of a custom recombination map. This gives users the flexibility to define their own recombination patterns according to specific genetic landscapes or experimental setups. This feature enables more realistic and tailored simulations that reflect the specific recombination patterns observed or desired in a study system.

#### Documentation

STUN is designed for a wide range of users, including those with little programming expertise, as well as advanced programmers interested in developing new fitness landscape models or statistics of quantitative genetics. To ensure a user-friendly experience, STUN provides a comprehensive manual within the package. The manual offers detailed explanations and guidance on all basic aspects and settings of the program. For users who wish to further explore the underlying model, implementation, and inner workings of STUN, code documentation is available. By running the command `cargo doc --open` in the terminal, the code documentation can be accessed and viewed in the default browser. This documentation provides additional details and explanations related to the code structure and functionality. The code documentation is recommended for users who intend to modify the code, require detailed information, or seek a deeper understanding of how the program operates.

### 2.3 Detailed output

One strength of STUN is that it can provide highly detailed information about the population and adaptation process. In addition to recording the genotype composition of the population in each generation or at defined time points, STUN offers a customizable suite of statistics that can be utilized to capture various aspects of the adaptive process, *e*.*g*., mean population fitness, haplotype diversity, fixation time of each allele, and population fitness distance to the global peak. By means of configuration files, users can examine the focal statistics at their discretion, hereby minimizing output files and saving efforts of later data processing. The frequency at which population information is recorded can be adjusted by setting the parameter *n*. For instance, if *n* = 1, population information will be collected and stored in every generation throughout the entire adaptive trajectory. Alternatively, users can choose a larger value of *n* to record data at regular intervals, providing a more aggregated view of the population dynamics. This allows users to strike a balance between data resolution and storage requirements. Evidently, users can output the full population data at any given generation of interest. This feature allows for a detailed examination of population features, individual genotypes and allele frequencies at specific time points. Additionally, at the end of each adaptive process, summary statistics are saved, which offer a concise overview of the evolutionary outcome. For further guidance on data collection, available statistics, and using the simulator effectively, users can refer to the user manual.

### 2.4 Performance

The performance of STUN is significantly influenced by the sizes of the fitness landscapes and the evolving populations. As both the fitness landscape and population increase in size, the mean computation time increases. To assess this relationship, we calculated the mean real time (wall clock time) for each simulation run, where 10 replicate populations sequentially adapted on a single fitness landscape using either standing genetic variation or new mutations (Figure 2). Remarkably, even with large sizes, such as a fitness landscape with 10 loci and a population of 10^4^ individuals, the computation time for 10 simulations remains relatively short. For instance, simulations involving standing genetic variation mostly require less than 2 seconds per simulation run. These results demonstrate STUN’s efficient handling of simulations, which makes it capable of managing large-scale simulations effectively.

**Figure 2.**
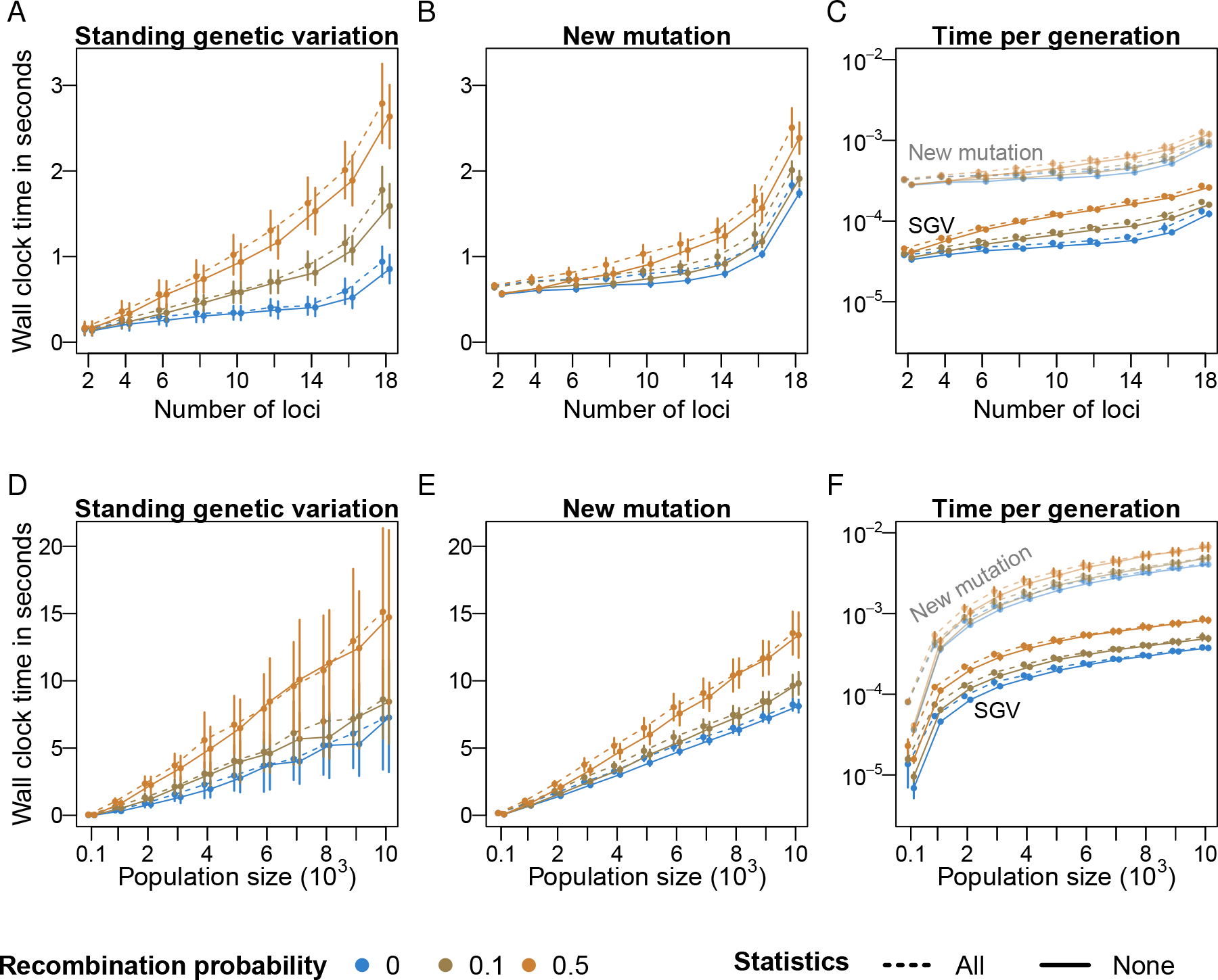
STUN performance indicated by wall-clock time in seconds for increasing numbers of loci and population size. We simulated adaptation on 100 landscapes and 10 populations for each fitness landscape. We recorded the real time for 10 simulations of a population adapting on a single landscape either using standing genetic variation (SGV) or using new mutations at a mutation rate of 1 *×* 10^*−*5^ per locus per generation (panels A, B, D and E). With standing genetic variation, the genetic drift threshold was set to 3, and the population size was 1000. For new mutations, the maximum generation number was 2000. Dots represent the mean of the real time, and bars represent one standard deviation of the real time. All simulations were performed on Rough Mount Fuji fitness landscapes, with additive and epistatic effects drawn from normal distributions *𝒩* (0, 0.01). Performance was measured in two conditions, either including the complete set of statistics available in the package (dashed lines) or none (solid lines). The average wall-clock time in seconds per generation is presented in panels C&F. All simulations were run on a 3.8 GHz Intel Core i7 CPU.

The performance of STUN depends differently on the number of loci for standing genetic variation (SGV) and new mutations. Regarding SGV, the computation time per generation scales linearly with the number of loci, despite the exponential growth of the number of genotypes of the greater underlying fitness landscape (Figure 2 A-C). With new mutations, as the number of loci increases, the mean wall-time dramatically increases (Figure 2B and 2C). This is explained by the increasing polymorphism resulting from new mutations, which leads to a large number of genotypes that segregate in the population at any time. Unlike the size of the fitness landscape, the population size influences computation time similarly for SGV and new mutations (Figure 2D & 2E), except for the variance. At high recombination rates and large population sizes (Figure 2D), the variance of wall time is considerable because in some simulations, several genotypes of similar fitness segregate for a long time (see 2.5; Figure 3). Overall, we hope that our presentation of the performance of STUN under different scenarios aids researchers in optimizing simulation settings for efficient and accurate simulations of complex trait adaptation.

**Figure 3.**
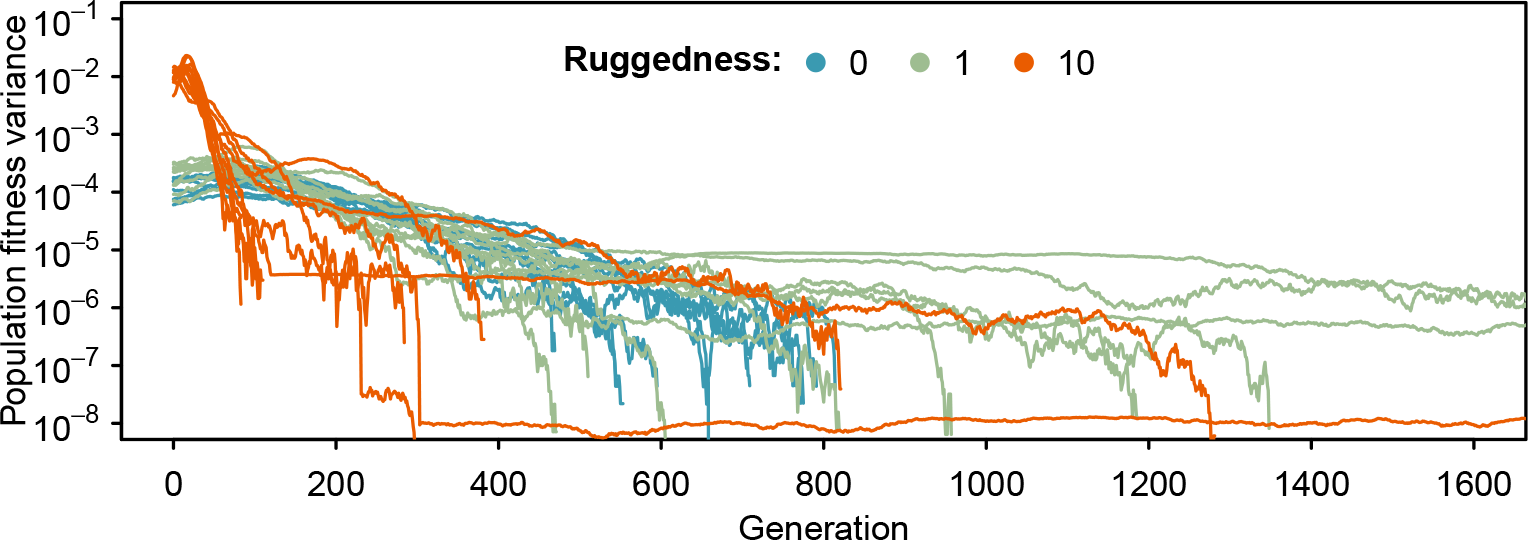
Dynamics of population fitness variance at different ruggedness levels. We simulated recombining populations adapting on Rough Mount Fuji fitness landscapes. The additive effect is constant (0.01), and epistasis is drawn from a normal distribution *N* (0, *σ*^2^). The ratio of the standard deviation (*σ*) of the epistasis parameter and the additive effect quantifies the ruggedness of the fitness landscape. Three ruggedness levels were considered in this example, 0, 1, and 10, where *σ* was 0, 0.01, and 0.1. We generated 10 fitness landscapes with 15 loci for each ruggedness level and let 1 population of size 5000 adapt to each landscape. The recombination rate is 0.1. The figure shows that epistasis can maintain segregating polymorphism for a long time.

### 2.5 Example analyses

In a previous study (Li et al. 2023), we used STUN to investigate how the interplay between recombination and epistasis affects adaptation. We found that the population follows a smoother-than-average adaptive path due to a higher marginal fitness of common genotypes. To further exemplify the practical utility of our program, we present an additional illustrative example. Specifically, we conducted simulations in which populations adapted to Rough Mount Fuji fitness landscapes with three levels of epistasis. Adaptation is assumed to happen from standing genetic variation. By tracking the variance in population fitness across generations, we demonstrate that a higher level of epistasis results in the rapid fixation of beneficial alleles (see Figure 3). Overall, this observation aligns with Fisher’s fundamental theorem of natural selection, as the presence of substantial fitness variance induced by epistasis amplifies the rate of adaptation, leading to shorter fixation times. On the other hand, it is noteworthy that some of our simulations exhibited a long-term non-zero variance in the population, indicating that multiple advantageous alleles coexist within the population. This suggests that epistatic interactions among loci may contribute to the maintenance of balanced polymorphism.

## 3 Limitations

In the current version of STUN, we focus on haploid recombining populations, which allows us to capture the relative effects of epistasis and additivity on the dynamics of adaptation on rugged fitness landscapes. However, we acknowledge that another type of non-additive effect, dominance in diploid populations, is not explicitly considered in STUN. In this context, it was proposed that dominance in diploid populations presumably could arise from epistatic interactions between alleles at the focal locus and other alleles at dominance modifier loci (*e*.*g*., Fisher 1928, Wright 1934, Kacser and Burns 1981, Wade 2001), but we are not aware of clear expectations or resulting fitness landscape models. Once the dominance distribution and its impact on the ruggedness of fitness landscapes has been documented empirically or proposed theoretically, incorporating dominance and diploid populations into STUN should be a worthy addition.

## 4 Conclusions

STUN enables the efficient simulation of rapid adaptation on rugged fitness landscapes, offering a comprehensive understanding of the complex dynamics involved in the adaptation of recombining organisms. By incorporating recombination and standing genetic variation, STUN captures the intricate interplay of genetic processes during the adaptive process. Moreover, STUN facilitates the utilization of existing empirical and theoretical fitness landscapes in simulations, allowing users to generate expectations regarding the complex evolutionary dynamics at play. One notable application of STUN is for the design of evolve-and-resequence experiments, where researchers aim to understand the genetic basis of adaptation by subjecting populations to controlled selection pressures. Using STUN, researchers could simulate and predict the dynamics of genetic variation, allele frequencies, and fitness trajectories in response to different environmental conditions. This information could help optimize experimental design, such as determining the appropriate selection strengths, population sizes, and sampling times, to achieve specific research objectives. Furthermore, STUN can provide a comprehensive understanding of the maintenance, elimination, or fixation of different types of mutations in evolving populations. By simulating the adaptive process, users can examine how different genetic factors, such as mutation rates, additive selection coefficients, and epistatic interactions, contribute to the rate and outcome of adaptation.

## Supporting information

Software manual

## Acknowledgements

We thank the THEE lab members for suggestions that improved this work. We acknowledge editorial suggestions from ChatGPT.

