## Supplementary material for "STUN: forward-time Simulation on TUnable fitNess landscapes in recombining populations": Software manual

### STUN: forward-time **S**imulation on **TU**nable fit**N**ess landscapes in recombining populations

This manual is provided as Supplementary material to STUN manuscript. A preprint can be found at https://doi.org/10.1101/2023.02.15.528681.

### 1 Overview

STUN simulates rapid adaptation on tunable fitness landscapes. The software integrates direct selection, epistasis, and recombination, and uses standing genetic variation or new mutations as the source of selected variants. STUN simulates the adaptation of haploid populations with recombination evolving on arbitrary fitness landscapes (Figure 1). In the initialization step, a population is generated by assigning the same allele frequency to the segregating alleles at each locus or by drawing from a folded neutral site frequency spectrum (SFS, Hudson [2015]; 4.1). When using the neutral SFS, it is possible to set a genetic drift threshold to exclude low-frequency alleles (Li et al. [2023]). The reproduction step is simulated by a Wright-Fisher model with non-overlapping, discrete generations. Within each generation, the individuals are randomly paired and perform recombination to generate a gamete pool. The recombination map can be uniform by assuming that the recombination probability is the same between neighboring loci (4.2.1). Alternatively, any custom recombination map can be used. Then, offspring are sampled from the gamete pool with a probability proportional to their fitness on the underlying fitness landscape to form the new generation (4.2.3). This process continues until all loci are monomorphic. In addition to generating the fitness landscapes and simulating the adaptation process, STUN provides a suite of customizable statistics (5).

#### STUN flow chart

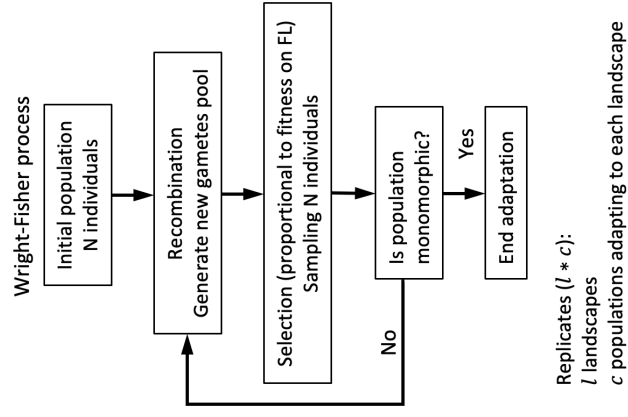

#### Modeling rapid adaption in recombining populations

1. Initialize neutral recombining population with $N$ haploid individuals
2. A sudden environmental change creates rugged fitness landscape (FL)
3. Evolution towards a monomorphic population or until a specific generation

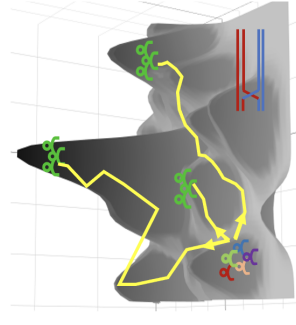

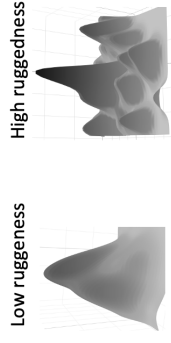

a) Mutations: mutation rate

b) Site frequency spectrum (SFS):
Uniform or Neutral SFS

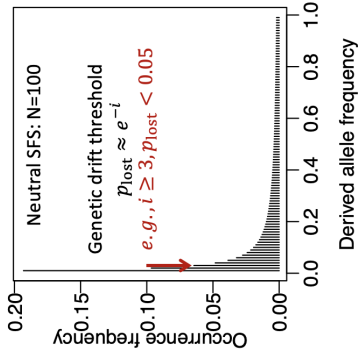

Fitness landscape models:

▪ Additive
▪ House of cards
▪ Rough Mount Fuji
▪ NK
▪ Block
▪ Custom...

Output:

• Population statistics
  ✓ Final genotype
  ✓ Fixation time
  ✓ Mean fitness of the population
  ✓ Fitness variance
  ✓ .....
• Raw population data
• Fitness landscape

Figure 1: Overview of STUN. STUN is a population-genetic simulation package designed for understanding complex trait adaptation. It employs Wright-Fisher dynamics and incorporates rugged fitness landscapes. STUN also offers a range of statistics on population features.

#### 2 Installation

We provide precompiled software executables for Linux, macOS and Windows (https://bit.ly/stun_manual_binaries).

*Note:* (macOS) If macOS blocks the executable from running due to security concerns, the user should compile the program using the instructions below.

*Note:* (Linux) If the executable does not have “execute” permission after download, the user should set this permission with the command `chmod +x stun_linux` before running STUN.

As an alternative, users can compile the software on a computer by following these steps:

1) Install rust toolchain. `stun` is written in rust. Instructions on how to install the latest rust toolchain can be found at https://www.rust-lang.org/tools/install

2) Clone the github repository https://github.com/banklab/STUN (or download and unzip it). Users can clone the repository with the instruction

```
git clone https://github.com/banklab/STUN.git
```

3) Compile the program. Use `cargo` (automatically installed with rust toolchain) to compile the program

```
cargo build --release
```

This creates the program `stun`. Notice that, by default, `cargo` will generate these binaries in the folder `target/release`.

4) (optional step, Linux or macOS) Add the program to the PATH. To call this program directly from anywhere in the command line, add the program to the environment parameter, PATH.

```
stun='pwd'/target/release
export PATH=$stun:$PATH
```

The `'pwd'` command writes the absolute pathname of the current working directory to the standard output. If your working directory is the folder compiling `stun`, “`stun='pwd'/target/release`” works. Otherwise, provide the absolute pathname containing the program.

5) (optional step) Generate code documentation. An automatic HTML version of the rust code documentation can be generated by `cargo` with the command

```
cargo doc
```

To open the code documentation in the default browser, run `cargo doc --open`. This code documentation is especially relevant if you plan to modify the code and need detailed information regarding the implementation or want to become familiar with the program's inner workings.

##### 3 Running a simulation

To demonstrate how to run STUN (`stun help`), we first give a brief example and then explain the parameter details in full in section 4.

In this example, we illustrate how to run a simulation. We consider a population with 200 individuals (`-N 200`). We assume that the initial population is in mutation-drift balance, such that the initial allele frequency in the population follows a neutral site frequency spectrum (`--neutralsfs`). We further assume that the population performs recombination with three recombination probabilities (0.001,0.01,0.1). To test the effect of recombination on adaptation, we impose three recombination probabilities (`--recombination_rates 0.001,0.01,0.1`). Selection is imposed on populations by means of a fitness landscape. In this example, we generate five () Rough Mount Fuji fitness landscapes, each with 10 loci (`-L 10`). In each fitness landscape, the alleles at each locus are coded as 0 or 1. Each allele coded 1 is assigned a constant additive effect $\mu_a = 0.01$ (`--mu_a 0.01`) and each genotype is assigned an epistatic coefficient drawn from a normal distribution with standard deviation $\sigma_e = 0.1$ (`--sigma_e 0.01`). For details of the Rough Mount Fuji model check section 4.2.6. With each set of parameters (parameters of recombination and selection), 50 replicate populations are simulated on each fitness landscape (`--replicates 50`).

```
stun -l 5 -N 200 --neutralsfs --recombination_rates 0.001,0.01,0.1 --replicates
50 RMF -L 10 --mu_a 0.01 --sigma_a 0 --sigma_e 0.01
```

The structure of this command is `stun <options> <model>`. *Note:* the options (in orange) should be declared first and then the model with its parameters (in purple). The model parameters are programmed as the secondary options.

The command above provides us with the default output. In the subsections below, we provide detailed descriptions of all options (4) and the customized output (5) of the program.

Note that the user can generate fitness landscapes without running evolutionary simulations with the command

```
stun -l 5 --generate_only RMF -L 10 --mu_a 0.01 --sigma_a 0 --sigma_e 0.01
```

Here, the `--generate_only` option signals that no simulations should be run.

#### 4 STUN: parameters and models

##### 4.1 Initializing a population

To initialize a population, the user specifies the population size and genotype composition (Figure 1). The population size is set using the option `--size <population_size>` (or the option `-N <population_size>` in short), where the `<population_size>` is an integer, the number of individuals in the population (1000 by default).

In the initial population, each individual is represented by one genotype. To obtain those genotypes, we assume that each locus is biallelic with minor (less frequent) and major (more frequent) alleles, where alleles are scattered randomly to form a group of genotypes in approximate linkage equilibrium. The minor allele frequency can be drawn from a folded neutral site frequency spectrum (Hudson [2015]) with or without a genetic drift threshold (see Li et al. [2023]), accessed through the option `--neutralsfs` (or `-n`). This option has two optional values: a drift threshold ($d$, integer value) can be specified as a first argument indicating the expected minimal copies of minor alleles in the neutral SFS to avoid alleles at a very low frequency in the initial population, $d = 1$ (i.e., no drift threshold) by default; and the probability ($p$, float value) that the minor allele is encoded as 1 as a second argument because allele 1 is presumed to carry the additive effect in some fitness landscape models, $p = 0.5$ by default.

The user can also set all alternative alleles to the same frequency. The user can choose the uniform model for the initial population with the option `--uniform <f>` (or `-u <f>`). In this model, the user specifies a probability $f$ that an allele is set as the alternative allele (encoded with a 1 in the code). And the allele frequencies at one locus are expected to be $f$ and $1 - f$ for allele 1 and 0, respectively. If $f = 0.5$, allele 1 and 0 are equally probable, meaning that the genotypes in the initial population are chosen at random with uniform probability.

In addition to standing genetic variation, a population can be initialized to be monomorphic by defining one random genotype (`--monomorphic random`) or any genotype on the fitness landscape (`--monomorphic <sequence>`). The polymorphism will be introduced by new mutations at a mutation probability per locus per generation (`-m`, `--mutation_probability <mutation_probability>`). The simulation will run until the defined maximum generation (`-t`, `--max_generation <max_generation>`).

#### 4.2 Adaptation on a fitness landscape: Wright-Fisher dynamics

Populations in STUN evolve according to the Wright-Fisher model of evolution with discrete non-overlapping generations (Figure 1). Each generation, the population performs recombination (4.2.1) and mutation (4.2.2) to generate a gamete pool. Then, selection happens as $N$ genotypes are sampled from this gamete pool with a probability proportional to the genotype fitness in the fitness landscape (4.2.3). The simulation runs until all polymorphism has been exhausted (if mutation probability is set to zero) or the maximum specified time has been reached.

##### 4.2.1 Recombination

Users of STUN can specify the recombination map in two ways.

If the map is uniform, i.e., the probability of recombination is the same between any pair of consecutive loci, the option `--recombination_rates` (or `-r`) is available. This option is followed by one or more recombination probabilities, separated by a comma, e.g., `-r 0.1` or `-r 0.01,0.1,0.5`. By default, `-r 0`, no recombination occurs.

The user can also specify arbitrary recombination maps, defining the recombination probability between all consecutive loci. This can be achieved with the option `--recombination_map`. This option takes either a list of recombination probabilities ($L - 1$ comma-separated integers) or the name of a file containing a list of recombination maps. Each recombination map should occupy one line in the file, and the values should be separated by a comma. Empty lines and comment lines starting with a `#` are allowed and ignored. Example 1 illustrates a possible recombination map file.

```
0.1,0.1,0.5,0.1
0,0.1,0.01,0.4

# each locus on a different chromosome
0.5,0.5,0.5,0.5
```

Example 1: File describing 3 recombination maps for a fitness landscape with 5 loci.

###### 4.2.2 Mutation

Mutation can be specified using the option `--mutation_probability <mutation_probability>` (`-m <mutation_probability>`). This mutation probability is a mutation probability per locus per generation, as such it should be a real number in the 0 to 1 range. If the mutation rate is large enough, several loci of a given individual in the population may mutate simultaneously.

###### 4.2.3 Selection: specifying the fitness model

Selection in STUN happen via an explicit genotype-fitness map, a so-called fitness landscape. We consider $L$ biallelic loci in the fitness landscape and alleles 0 and 1 at each locus ($g_i \in \{0, 1\}$). The genotypes are $\vec{g} = [g_1, g_2, \dots, g_L]$. The fitness of a genotype $\vec{g}$ is referred to as $f(\vec{g})$, which corresponds to its selection strength in the polymorphic population. We implement five fitness landscape models (additive, House of Cards, Rough Mount Fuji, NK, and Block models). The Rough Mount Fuji, NK, and Block models contain a tunable amount of epistasis, quantified by their ruggedness. Additive and House of Cards models are limiting cases of no and maximum epistasis in the other models. All models are described in the following.

In STUN, fitness landscapes are specified either using one of the five out-of-the-box model implementations or arbitrarily defined in a custom file. The five models are described in the section “Models” in `stun help`. The model parameters can also be obtained by specifying a model's name after `stun help`. For example, for checking the detailed options for the Rough Mount Fuji model, run the command

```
stun help RMF
```

The fitness landscape models are implemented as subcommands of STUN and should be specified *always after all remaining options*. In the following subsections, we provide detailed descriptions of the five implemented models and how to use custom fitness landscapes.

###### 4.2.4 Additive model

The additive model assumes that allele 0 is the reference allele with fitness 0 and assigns an additive fitness effect to each allele 1. Additive effects are drawn from a normal distribution $\mathcal{N}(\mu, \sigma^2)$, with mean $\mu$ and standard deviation $\sigma$. Thus, the fitness of a genotype ($\vec{g}$) can be expressed as

$$f(\vec{g}) = \sum_{i=1}^L g_i a_i, \quad (1)$$

where $a_i$ is the fitness effect assigned to allele 1 at locus $i$.

The ruggedness of a fitness landscape describes the level of fitness contribution from epistasis (i.e., genetic interaction between loci for fitness) compared to that from the additive effect. Since no epistasis is assumed in the additive model, the ruggedness of the resulting fitness landscapes is always 0. The additive model is the no-epistasis limiting case of the Rough Mount Fuji model (setting the epistatic parameter to zero), NK model (setting the number of interacting loci $K$ to zero), and the Block model (setting the number of blocks equal to the number of loci).

The additive model is specified in STUN by the model option `additive` with three parameters,

• `-L <number_of_loci>` number of loci of the fitness landscape [default $L = 5$ loci].
• `-m <mu>` or `--mu <mu>` mean of the fitness effects [default $\mu = 0.1$].
• `-s <sigma>` or `--sigma <sigma>` standard deviation of the fitness effects [default $\sigma = 0.1$].

As an example, to generate an additive fitness landscape with 8 loci, a constant additive effect ($\mu = 0.01$, and $\sigma = 0$) at each locus, we would run the command

```
stun --generate_only additive -L 8 --mu 0.01 --sigma 0
```

Equivalently, a more compact version is

```
stun --generate_only additive -L 8 -m 0.01 -s 0
```

###### 4.2.5 House of Cards model

The House of Cards model assigns random statistically independent fitness values to each genotype. In our implementation, each genotype has a fitness drawn from a normal distribution

$\mathcal{N}(\mu, \sigma^2)$, with mean $\mu$ and standard deviation $\sigma$.

By assigning statistically independent fitness values to each genotype, the House of Cards model corresponds to the limit of maximal ruggedness of the Rough Mount Fuji model (setting the additive contribution to zero), NK model (setting the number of interacting loci $K$ to $N - 1$), or the Block model (setting the number of blocks to 1).

The House of Cards model is specified by the model option `HoC` with three options,

• `-L <number_of_loci>` number of loci of the fitness landscape [default $L = 5$ loci].
• `-m <mu>` or `--mu <mu>` mean fitness [default $\mu = 0.1$].
• `-s <sigma>` or `--sigma <sigma>` standard deviation of the fitness [default $\sigma = 0.1$].

For example, to generate 10 House of Cards fitness landscapes with 8 loci, $\mu = 0$, and $\sigma = 0.01$, run the command

```
stun --generate_only --landscapes 10 HoC -L 8 --mu 0 --sigma 0.01
```

Equivalently,

```
stun --generate_only -l 10 HoC -L 8 - 0 -s 0.01
```

###### 4.2.6 Rough Mount Fuji model

The Rough Mount Fuji model was introduced in Aita et al. [2000]. Fitness is defined by two terms: an additive term that assigns a fitness effect to allele 1 at each locus; and an epistatic term that is assigned to each genotype. The additive effects are drawn from a normal distribution $\mathcal{N}(\mu_a, \sigma_a^2)$, with mean $\mu_a$ and standard deviation $\sigma_a$. The epistatic effects are drawn from a normal distribution $\mathcal{N}(0, \sigma_e^2)$, with zero mean and standard deviation $\sigma_e$. Mathematically, the fitness of a genotype on the Rough Mount Fuji fitness landscape is

$$f(\vec{g}) = \sum_{i=1}^L g_i a_i + \epsilon_{\vec{g}}, \quad (2)$$

where $a_i$ is the additive effect at locus $i$ and $\epsilon_{\vec{g}}$ is the epistatic effect of genotype $\vec{g}$.

When additive effects are equal for all loci (i.e., $\sigma_a = 0$), the ratio between the epistatic standard variation $\sigma_e$ and the additive effect $\mu_a$ describes the ruggedness of the fitness landscape. A higher ratio indicates a higher ruggedness.

The Rough Mount Fuji model is specified by the model option `RMF` with four parameters,

• `-L <number_of_loci>` number of loci of the fitness landscape [default $L = 5$ loci].

• `-m <mu_a>` or `--mu_a <mu_a>` mean additive fitness effects [default $\mu_a = 0.1$].
• `-s <sigma_a>` or `--sigma_a <sigma_a>` standard deviation of the additive effects [default $\sigma_a = 0.1$].
• `-S <sigma_e>` or `--sigma_e <sigma_e>` standard deviation of the epistatic effects [default $\sigma_e = 0.1$].

For example, to generate 10 Rough Mount Fuji fitness landscapes with 8 loci, $\mu_a = 0$, $\sigma_a = 0.01$, and $\sigma_e = 0.02$, run the command

```
stun --generate_only --landscapes 10 RMF -L 8 --mu_a 0 --sigma_a 0.01 --sigma_e 0.02
```

or, equivalently,

```
stun --generate_only -l 10 RMF -L 8 -m 0 -s 0.01 -S 0.02
```

###### 4.2.7 NK model

The NK model was proposed by Kauffman and Levin [1987]. In the NK model, $N$ is the number of loci, and $K$ is the number of loci that a focal locus interacts with. $K$ can vary from 0 to $N - 1$. Larger $K$ implies a larger ruggedness because the contribution of each locus depends on more loci. One limiting case is $K = 0$, where no interactions occur between loci. This limiting case corresponds to an additive landscape (see 4.2.4). The other limiting case is $K = N - 1$, meaning that the fitness of each locus depends on its interactions with all other loci, which results in a House of Cards fitness landscape (see 4.2.5). Intermediate $K$ values span the ruggedness between these two extreme cases.

The NK model is specified by the model option `NK` with two parameters

• `-N <number_of_loci>` number of loci of the fitness landscape [default $N = 5$ loci].
• `-K <K>` number of interacting loci per focal locus. $K$ is constrained to the range $[0, L - 1]$ [default $K = 0$ loci].

For example, for generating 10 NK fitness landscapes with $N = 8$ loci, each locus interacting with $K = 3$ loci, run the command

```
stun --generate_only -l 10 NK -N 8 -K 3
```

###### 4.2.8 Block model

The Block model was introduced in Perelson and Macken [1995] and assumes that epistatic interactions happen only among loci within a block. Consider $L$ loci on the fitness landscape. $L$

loci are randomly assigned to $B$ blocks. All blocks are independent of each other. Within each block, the fitness is determined by a normal distribution, i.e., each block behaves like a small House of Cards model. In STUN, the epistatic coefficients are drawn from a normal distribution $\mathcal{N}(\mu, \sigma^2)$, with mean $\mu$ and standard deviation $\sigma$.

The Block model is a relative of the NK model. Similarly to the NK model, ruggedness is determined by the number of blocks, which vary from 1 to $L$. A Block model with $L$ blocks is equivalent to the additive model (see 4.2.4); block models with 1 block represent the House of Cards model (see 4.2.5). Any intermediate number of blocks indicates intermediate ruggedness.

The Block model is specified by the model option `block` with four parameters

• `-N <number_of_loci>` number of loci of the fitness landscape [default $N = 5$ loci].
• `-B <number_of_blocks>` number of blocks (in the range of 1 to $N$) [default $B = 1$ block].
• `-m <mu>` or `--mu <mu>` mean epistatic coefficient of genotypes within a block [default $\mu = 0$].
• `-s <sigma>` or `--sigma <sigma>` standard deviation of the epistatic coefficients within a block [default $\sigma = 0.1$].

For example, to generate 10 fitness landscapes according to the Block model, with 8 loci and 3 blocks each, and within each block, with $\mu = 0$ and $\sigma = 0.01$, run the command

```
stun --generate_only -l 10 block -N 8 -B 3 --mu 0 --sigma 0.01
```

or

```
stun --generate_only -l 10 block -N 8 -B 3 -m 0 -s 0.01
```

###### 4.2.9 Custom fitness landscapes

In addition, STUN allows for the definition of arbitrary fitness landscapes. A provided fitness landscape file has to consist of a list of genotypes and their respective fitness values. Each line should specify a genotype with $L$ biallelic alleles (coded as 0 or 1) and a fitness value for the genotype, separated by any type of ASCII white space. The genotypes can appear in any order. *Any lines starting with the character # will automatically be ignored.* This format is illustrated in Example 2.

Biallelic fitness landscapes generated by Magellan (Brouillet et al. [2015]) are also accepted as input, https://bioinfo.mnhn.fr/abi/public/Magellan/Magellan.main.html. Magellan's website and program provide tools for generating a wide diversity of fitness landscapes as well as a curated list of experimental fitness landscapes which can be directly used as input to our program. Whereas Magellan accommodates multiallelic fitness landscapes, STUN is restricted to biallelic fitness landscapes.

For example, a fitness landscape file specifying a three-locus fitness landscape has the following structure:

```
# made-up fitness landscape
0 0 0 0.7231
1 1 1 0.7312
0 0 1 0.2731
0 1 1 0.1327
1 1 1 0.2317
0 1 0 0.7321
# next genotype is very important
1 1 0 0.1723
1 0 0 0.1732
```

Example 2: File that specifies a custom fitness landscape.

*Note:* 1. lines starting with # are ignored; 2. genotypes appear in an arbitrary order.

Option `custom --file fl.dat` loads a fitness landscape file. For example, to simulate a population initially under mutation-drift balance (`--neutralsfs`) that adapts on the fitness landscape described in the file `fl.dat`, run the command

```
stun --neutralsfs custom --file fl.dat
```

or, equivalently,

```
stun -n custom -f fl.dat
```

STUN can also load a group of fitness landscape files. The first way is using `custom --file` with asterisks (\*) in the file name used as a wildcard. All files matching the pattern will be used by STUN. For example, if the user wants to use three fitness landscape files in a folder `fitness_landscapes`, `fl1.dat`, `fl2.dat` and `fl55.dat`, they should run the command

```
stun -n custom -f fitness_landscapes/fl\*.dat
```

STUN will load all three fitness landscapes. *Note:* the backslash before the \* is necessary for most command lines because the command line will automatically match the wildcard when the backslash is missing, before passing the instruction to STUN.

The second way to use a group of fitness landscapes is the option `--list`. This option is followed by a file to specify a list of fitness landscapes (see Example 3). This list file should contain the path of each fitness landscape, one fitness landscape path per line. Empty and comment lines (starting with a #) are ignored. In this list, the order of fitness landscapes matters because the order in the list file determines the indices of landscapes in the output (with 0 being the first landscape, 1 the second, ...). An example list file looks as follows:

```
# list of fitness landscapes
landscapes/example_fitness_landscape1.dat
landscapes/example_fitness_landscape2.dat

landscapes/another_example_fitness_landscape.dat
```

Example 3: File listing paths of fitness landscapes.

#### 5 Output

STUN has three types of outputs (Figure 1) – statistics about population features (Section 5.1), raw population data (Section 5.2), and fitness landscapes (Section 5.3). The user can customize the output, choosing from a wide range of options available. Choosing a smaller set of options may result in a better running time because only the options specified by the user are calculated (Section ??). Custom output is specified in a configuration file (option `--output_config <output_configuration_file>`, where `<output_configuration_file>` is the configuration file name), written in a simple configuration file format, TOML<sup>1</sup>. We provide a file including all options, `example_output_configuration.toml`, which can be found at the root of the program repository. To match the user's needs, the output can be easily configured with simple modifications to this file, e.g., by commenting/uncommenting lines. Following is the explanation of each term in the configure file (Sections 5.1 and 5.2).

##### 5.1 Population statistics

###### 5.1.1 Periodic statistics

The first section in the file consists of population statistics which can be saved periodically. Example 4 shows a possible configuration for this section.

<sup>1</sup>https://toml.io/

```
#####
# Saves periodic information about the population
[details]
# set to true to save periodic information
save = true

# uncomment to specify the file name (otherwise an automatic file name
# is assigned)
# output_filename = "example_%N.dat"

# save the data every 'period' generations (default: 1)
period = 1

# list of possible options to save, comment or uncomment options
options = [
    "mean_fitness",
    "variance_fitness",
    # "median_fitness",
    # "maximum_fitness",
    # "minimum_fitness",
    # "median_fitness_with_genotype",
    # "maximum_fitness_with_genotype",
    # "minimum_fitness_with_genotype",
    # "shannon_entropy",
    "haplotype_diversity",
    # "genotypes_count",
    # "major_genotype",
    "fixations_count"
]
#####
```

Example 4: Output configuration file section regarding the periodic population information.

In the example 4, the section starts with the key [details]. Setting save to false will cause the program to ignore all periodic statistics. When save=true, period sets the saving period of the statistics, e.g., period = 10 means saving the statistics every ten generations; output\_filename allows the user to override the default output file name<sup>2</sup> (see Box 1 for more information on output file names); options specifies which statistics to calculate and save.

The output file is a tab-separated file, saved in the folder data/population\_details. The first line contains a comment with the column names. The first four columns are reserved for the global parameters of the simulation run, ordered as the landscape id, the recombination rate, the replicate id, and the generation number. The following columns contain the statistics requested by the user and in order of their appearance in the configuration file.

<sup>2</sup>Notice the %N in the filename in the example. %N is automatically replaced by the population size in the actual name of the file. A full list of replacements can be found in Box 1.

The full list of statistics available is

• Mean fitness of the population ("mean\_fitness")
• Fitness variance in the population ("variance\_fitness")
• Median of population fitness, with the option of saving also the corresponding genotype (respectively, "median\_fitness" and "median\_fitness\_with\_genotype")
• Minimum fitness in the population, with the option of saving also the corresponding genotype (respectively, "minimum\_fitness" and "minimum\_fitness\_with\_genotype")
• Maximum fitness in the population, with the option of saving also the corresponding genotype (respectively, "maximum\_fitness" and "maximum\_fitness\_with\_genotype")
• Shannon diversity of the population ("shannon\_diversity")
• Haplotype diversity ($1 - \sum_g p_g$, where $p_g$ is the frequency of the genotype $g$) ("haplotype\_diversity")
• Genotype count, i.e., the number of genotypes present in the population ("genotypes\_count")
• Major genotype, i.e., the composite genotype that has all alleles as the major alleles in the population ("major\_genotype")
• Fixation count, i.e., the number of fixations that occurred until the current generation ("fixations\_count")

##### 5.1.2 Fixation details

The user can save information about the fixation steps that occur during the adaptation process. With this option, a list including all fixation events will be logged to a file. Notice that, when mutation is allowed a fixed mutation can become polymorphic again, so each allele can undergo several fixation events. This output can be activated/deactivated with the variable `save` in the section `[fixation_details]` of the configuration file, as shown in Example 5. The output file name can be customized by defining a variable `output_filename` in this section. Example 6 shows an excerpt of a fixation details output file with the landscape id, recombination rate, mutation probability and replicate index occupying the first four columns, followed by the fixation time, the locus and the fixed allele.

All output file names can use a set of replacement characters that generate the file names dynamically, e.g., "RMF\_%L\_N%N.dat" will be rendered as RMF\_5\_N1000.dat if the number of loci is 5 and the population size 1000. Here is the full list of replacements

• %m: model's name (e.g., RMF for Rough Mount Fuji model)
• %M: model description (e.g., RMF\_L5\_mu0\_stda0.1\_stdb0.1 for a Rough Mount Fuji model with 5 loci and corresponding parameters)
• %L: number of loci
• %N: population size
• %l: number of landscapes to use
• %id: user-specified identifier (by --id parameter in the command line)
• %I: initial population description (e.g., neutralsfs\_dt2\_p0.5 for a population initiated with a neutral SFS with drift threshold 2 and minor allele probability 0.5)
• %i: initial population in shorter form (only shows the name of initial population model, e.g., neutralsfs for the previously described situation)

*Note:* the default file name contains the model description "%M.dat"; if the user specifies an id, it appends it to the file name "%M\_%id.dat".

Box 1: Output file name replacements.

```
#####
# Saves information about each fixation in the population (saved whenever a
# fixation occurs)
[fixation_details]
# set to true to save fixation details
save = true

# uncomment to specify the file name (otherwise an automatic file name is
# assigned)
# output_filename = "example.dat"
#####
```

Example 5: Output configuration file section to log every fixation event.

```
#landscape_id recombination_rate mutation_probability replicate fixation_time
#locus allele
0 0.1 0.001 0 1 2 0
0 0.1 0.001 0 2 3 0
0 0.1 0.001 0 4 11 0
0 0.1 0.001 0 5 0 0
0 0.1 0.001 0 17 2 0
0 0.1 0.001 0 748 9 1
0 0.1 0.001 0 750 9 1
0 0.1 0.001 0 754 9 1
1 0.1 0.001 0 1 0 0
1 0.1 0.001 0 1 6 0
1 0.1 0.001 0 2 3 0
1 0.1 0.001 0 3 6 0
2 0.1 0.001 0 2 0 0
...
```

Example 6: Excerpt of a fixation details output file.

##### 5.1.3 Final statistics

At the end of each run, four features are saved by default, namely

• the number of generations until fixation of all loci
• whether the final genotype is the absolute fitness maximum on the underlying fitness landscape.
• the final fitness
• the adaptive load of the population, the fitness differences between the final genotype and the global peak normalized by the fitness of the global peak.

This output can be activated/deactivated with the flag `save` in the section `[final_statistics]` of the configuration file, as shown in Example 7. The output file name can be customized by defining a variable `output_filename` in this section. Example 8 shows an excerpt of a final statistics output file with the landscape id, recombination rate, mutation probability and replicate index occupying the first four columns, followed by the data.

```
#####
# Saves final statistics for each run
[final_statistics]
# set to true to save final statistics
save = true

# uncomment to specify the file name (otherwise an automatic file name
# is assigned)
# output_filename = "example.dat"
#####
```

Example 7: Output configuration file section regarding saving the final statistics.

```
#landscape_id recombination_rate mutation_probability replicate tfixation
#found_maximum adaptive_load final_fitness
0 0 0 0 51 1 0 0.7160064320989423
0 0 0 1 17 0 0.12670956185612903 0.6252815708015151
0 0 0 2 27 1 0 0.7160064320989423
0 0 0 3 25 0 0.18980510862560548 0.5801047534777704
...
```

Example 8: Excerpt of a final statistics output file.

#### 5.2 Raw population data

The detailed composition of the population can be saved at the initial time and periodically, as defined in the section [population] of the output configuration file. The flag `initial` saves the initial population when set to true. If the user also wishes to save the population periodically, they can set that with the variable `period`. Setting the period to 0 forces STUN to ignore the periodic saves. Note that saving the population too often may slow down the simulation or consume large amounts of disk space. Example 9 illustrates the options. Example 10 shows a possible output for an initial population adapting on a 3 loci fitness landscape. The first three columns identify the simulation run, whereas the others represent pairs of genotype and genotype counts. Only genotypes segregating in the population are represented, i.e., there are no genotypes with a zero count. Periodic population files follow a similar structure.

```
#####
# Saves population
[population]
# set to true to save the initial population
initial = true

# uncomment to specify the file name (otherwise an automatic file name
# is assigned)
# output_filename_initial_population = "example.dat"

# save population periodically (set to 0 to ignore)
# warning: saving large populations too often can slow down the
#          simulation and/or consume a large disk space
period = 0

# uncomment to specify the file name (otherwise an automatic file name
# is assigned)
# output_filename_periodic_population = "example.dat"
#####
```

Example 9: Output configuration file section regarding saving the full population.

```
#landscape_id recombination_rate mutation_probability replicate genotype count
...
0 0 0 0 000 3 100 1 010 3 110 3 001 288 101 109 011 436 111 157
0 0 0 1 100 3 010 3 110 988 111 6
0 0 0 2 100 18 110 2 001 6 101 922 111 52
0 0 0 3 000 823 100 32 010 135 110 3 001 5 011 2
...
```

Example 10: Excerpt of an initial population output file.

##### 5.3 Fitness landscapes

Fitness landscapes generated by the program are saved in the folder `landscapes/`. Two files are saved for each landscape: one with a `.bin.fl` extension and another with `.dat` extension. The `.bin.fl` file is a binary file representing the landscape used by STUN and can be ignored by the user. The `.dat` file is a human-readable file that starts with comments describing the model followed by a list of genotypes and their corresponding fitness. The `.dat` file is saved in the format described for the input of custom fitness landscapes (see Example 2). The first lines are always occupied by comments describing the model used, followed by the list of genotypes and corresponding fitnesses.

Example 11, which shows a NK fitness landscape with 6 loci and $K = 3$, was obtained with the command

```
stun --generate_only NK -N 6 -K 3
```

```
# NK fitness landscape
# N: 8
# K: 3
0 0 0 0 0 0 0 0 0.6372950288693936
1 0 0 0 0 0 0 0 0.6764385965949848
...
0 1 1 1 1 1 1 1 0.4390467543484846
1 1 1 1 1 1 1 1 0.4383673839878178
```

Example 11: Output file for a fitness landscape.

#### 6 Example

We provide a simple example to demonstrate the utility of the software, available in the folder `example`. Fisher's fundamental theorem of natural selection describes the rate of fitness increase depending on population fitness variance in an additive model (original edition published in 1930; Coop [2021]). We used a Rough Mount Fuji fitness landscape to show how epistasis affects the rate of fitness increase. The rate of fitness increase is an indicator of the efficiency of selection since the increase in population fitness is confined by the number of polymorphic loci and the strength of selection (direct selection and epistasis). In our example, the additive effect is constant (0.01), and epistasis is drawn from a normal distribution $N(0, \sigma^2)$. The ratio of the standard deviation of the epistasis parameter and the additive effect quantifies the ruggedness of the fitness landscape. Three ruggedness levels are considered in this example, 0, 1, and 10, where $\sigma$ are 0, 0.01, and 0.1. We generated 10 fitness landscapes with 15 loci for each ruggedness level and let 1 population of size 5000 adapt to each landscape. We plotted population fitness variance over time. High ruggedness (10) results in a much larger population fitness variance than low ruggedness (0 and 1). Moreover, at high ruggedness the variance decays quickly and fixation occurs most rapidly. In contrast, adaptation under weak epistasis (ruggedness equal to 1) does not display increased population fitness variance compared to additive fitness landscapes in most of the simulations. Therefore, weak epistasis does not speed up fixation much. Notably, some populations take a long time to become monomorphic in the presence of epistasis because epistasis can maintain more than one genotype in long-term competition, resulting in a transient signal of balancing selection.

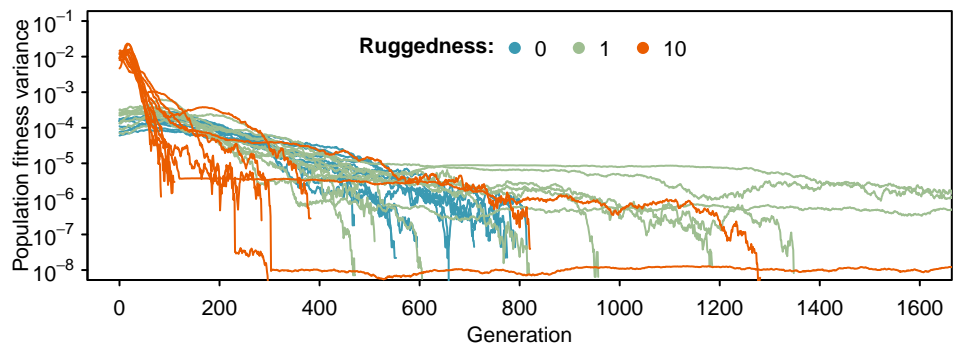

Figure 2: Dynamics of population fitness variance at different ruggedness levels.

#### 7 Additional information

Users are welcome to share the program freely and modify/adapt it for any purpose under an MIT License. Please cite our work if you use STUN https://doi.org/10.1101/2023.02.15.528681. For up-to-date contact information of the authors see https://banklab.github.io.
